# Furin cleavage of SARS-CoV-2 Spike promotes but is not essential for infection and cell-cell fusion

**DOI:** 10.1101/2020.08.13.243303

**Authors:** Guido Papa, Donna L. Mallery, Anna Albecka, Lawrence Welch, Jérôme Cattin-Ortolá, Jakub Luptak, David Paul, Harvey T. McMahon, Ian G. Goodfellow, Andrew Carter, Sean Munro, Leo C. James

**Author notes:** Correspondence (L.C.J.).

## Abstract

Severe acute respiratory syndrome coronavirus 2 (SARS-CoV-2) infects cells by binding to the host cell receptor Ace2 and undergoing virus-host membrane fusion. Fusion is triggered by the protease TMPRSS2, which processes the viral Spike (S) protein to reveal the fusion peptide. SARS-CoV-2 has evolved a multibasic site at the S1-S2 boundary, which is thought to be cleaved by furin in order to prime S protein for TMPRSS2 processing. Here we show that CRISPR-Cas9 knockout of furin reduces, but does not prevent, the production of infectious SARS-CoV-2 virus. Comparing S processing in furin knockout cells to multibasic site mutants reveals that while loss of furin substantially reduces S1-S2 cleavage it does not prevent it. SARS-CoV-2 S protein also mediates cell-cell fusion, potentially allowing virus to spread virion-independently. We show that loss of furin in either donor or acceptor cells reduces, but does not prevent, TMPRSS2-dependent cell-cell fusion, unlike mutation of the multibasic site that completely prevents syncytia formation. Our results show that while furin promotes both SARS-CoV-2 infectivity and cell-cell spread it is not essential, suggesting furin inhibitors will not prevent viral spread.

## INTRODUCTION

In late 2019, a new coronavirus was identified in a Chinese province, Hubei, and named severe acute respiratory syndrome coronavirus 2 (SARS-CoV-2) for its close similarity to severe acute respiratory syndrome (SARS-CoV-1) that appeared in 2002. SARS-CoV-2 spread rapidly, so that the World Health Organisation had to declare a pandemic on March 11^th^. By 13^th^ August, it had infected more than 21 million people and caused more than 750 thousand deaths worldwide. Several studies have shown that the introduction of the SARS-CoV-2 into humans was the result of zoonotic transmission, with pangolins acting as an intermediary host between bats and humans (Andersen et al., 2020; Lam et al., 2020).

SARS-CoV-2 (CoV-2) entry into host cells is mediated by the spike (S) protein, which is a major surface protein incorporated into the viral envelope (Shang et al., 2020). S protein is a trimeric transmembrane glycoprotein whose ectodomain includes two main subunits: S1, responsible for attachment to the host cell receptor Ace2 and for shielding the S2 subunit that contains the fusion machinery (Hoffmann et al., 2020a). Similar to the fusion proteins of many other respiratory viruses, S protein of SARS-CoV-2 is activated by cellular protease-mediated cleavage (Böttcher et al., 2006; Estes et al., 1981; Nagai et al., 1976; Scheid and Choppin, 1984). Activation of S requires proteolytic cleavage at two distinct sites: in the unique multi-basic site motif of Arg-Arg-Ala-Arg (RRAR), located between the S1 and S2 subunits, and within the S2 subunit (S2’) located immediately upstream of the hydrophobic fusion peptide that is responsible for triggering virus-cell membrane fusion (Madu et al., 2009; Millet and Whittaker, 2015; Walls et al., 2020).

CoV-2 S is a type I transmembrane protein which, besides being incorporated into virions and mediating viral entry, is present at the plasma membrane of infected cells. Plasma membrane-displayed S protein can trigger fusion with neighbouring cells, leading to the formation of enlarged, multinucleate enlarged cells (termed syncytia) (Buchrieser et al., 2020; Hoffmann et al., 2020b; Ou et al., 2020). This event is dependent upon expression of the cell-surface serine protease TMPRSS2, which exposes the S fusion peptide and makes S protein fusogenic directly at the host cell membrane (Buchrieser et al., 2020). These syncytia have also been found in tissues from individuals infected with SARS-CoV-2, suggesting an involvement in pathogenesis (Giacca et al., 2020; Tian et al., 2020; Xu et al., 2020).

One significant difference between S protein of SARS-CoV-2 and SARS-CoV-1 is that only the former contains a multi-basic cleavage site (Hoffmann et al., 2020b; Walls et al., 2020). The acquisition of a multibasic site by insertion of basic amino acids into surface glycoproteins has long been known to be a central virulence factor for many pathogenic viruses, including pandemic influenza viruses (Luczo et al., 2015; Schrauwen et al., 2012; Suguitan et al., 2012). This is thought to be because viruses with a multibasic site are readily cleaved by ubiquitously expressed proprotein convertases and can thus spread rapidly. In the case of SARS viruses, inhibitor experiments have suggested that the protease responsible for cleaving S protein is furin (Bestle et al., 2020; Hoffmann et al., 2020b). Furin is a calcium-dependent protease that recognise and cleaves the specific sequence motif R-X-R/K- R, where X is identified as any amino acid residue (Shiryaev et al., 2013; Walker et al., 1994). Furin is mainly expressed in the trans-Golgi network with little present in intracellular vesicles, suggesting that it may act on S protein during viral production (Hoffmann et al., 2020b; Izaguirre, 2019). In the context of SARS-CoV-2, the acquisition of a multibasic site in the S protein has been identified as a determinant of pandemic potential but its precise role in transmission remains unclear (Menachery et al., 2019; Yang et al., 2015). Here we sought to determine whether furin is essential for CoV-2 S multibasic site processing and how it impacts both viral entry and cell-cell spread.

## RESULTS

### CoV-2 S protein mediates cell-cell fusion between different cell types

Previous reports have shown that CoV-2 S protein possesses high fusogenic activity and is able to trigger large syncytia formation *in vitro* and *in vivo*, contrary to the S protein of two related coronaviruses SARS and MERS (Hoffmann et al., 2020b). Whether cell-cell fusion and formation of syncytia can occur between uninfected and infected cells and if it can happen between different cell types are significant questions in understanding CoV-2 pathogenesis. To address these questions, we established a CoV-2-S protein-mediated cell–cell fusion assay employing 293T cells stably overexpressing human ACE2 receptor (herein named 293T-hACE2) (Fig. S1A) and Vero cells, as both have been widely used in coronavirus studies (Kaye et al., 2006). Our assay is based on the ectopic overexpression of CoV-2 S together with the mCherry fluorescent protein in one cell type (named the donor cell) and the labelling of another cell type (named the ***acceptor* cell**) with a green fluorescent dye (Fig. 1A). Mixing both cell types and measuring the kinetics of merged fluorescence allows precise quantification of cell-cell fusion. Using this assay, we investigated the fusogenic properties of S protein when at the cell surface. We found that CoV-2 S-mediated syncytia formation occurred regardless of whether Vero or 293Ts were used as donor or acceptor cells (Fig. 1B,C). Moreover, we observed similar kinetics of syncytia formation whether we used Vero cells as both donor or acceptor or just one or the other. Next, we compared the impact of Ace2 overexpression with that of the protease TMPRSS2, which is responsible for cleavage and exposure of the fusion peptide prior to fusion. Importantly, overexpression of the TMPRSS2 protease in either Vero or 293T cells, markedly increased the rate of cell-cell fusion (Fig.1B,C,D, S1A). When TMPRSS2 was overexpressed on acceptor cells, syncytia formation reached almost 85% within 24 hours post transfection (Fig. 1D). This indicates that TMPRSS2 can be rate limiting for cell-cell fusion and highlights its fundamental importance in CoV-2 spreading. The data also suggest that the fusion machinery of SARS-CoV-2 is an important target for development of coronavirus antivirals.

**Figure 1.**
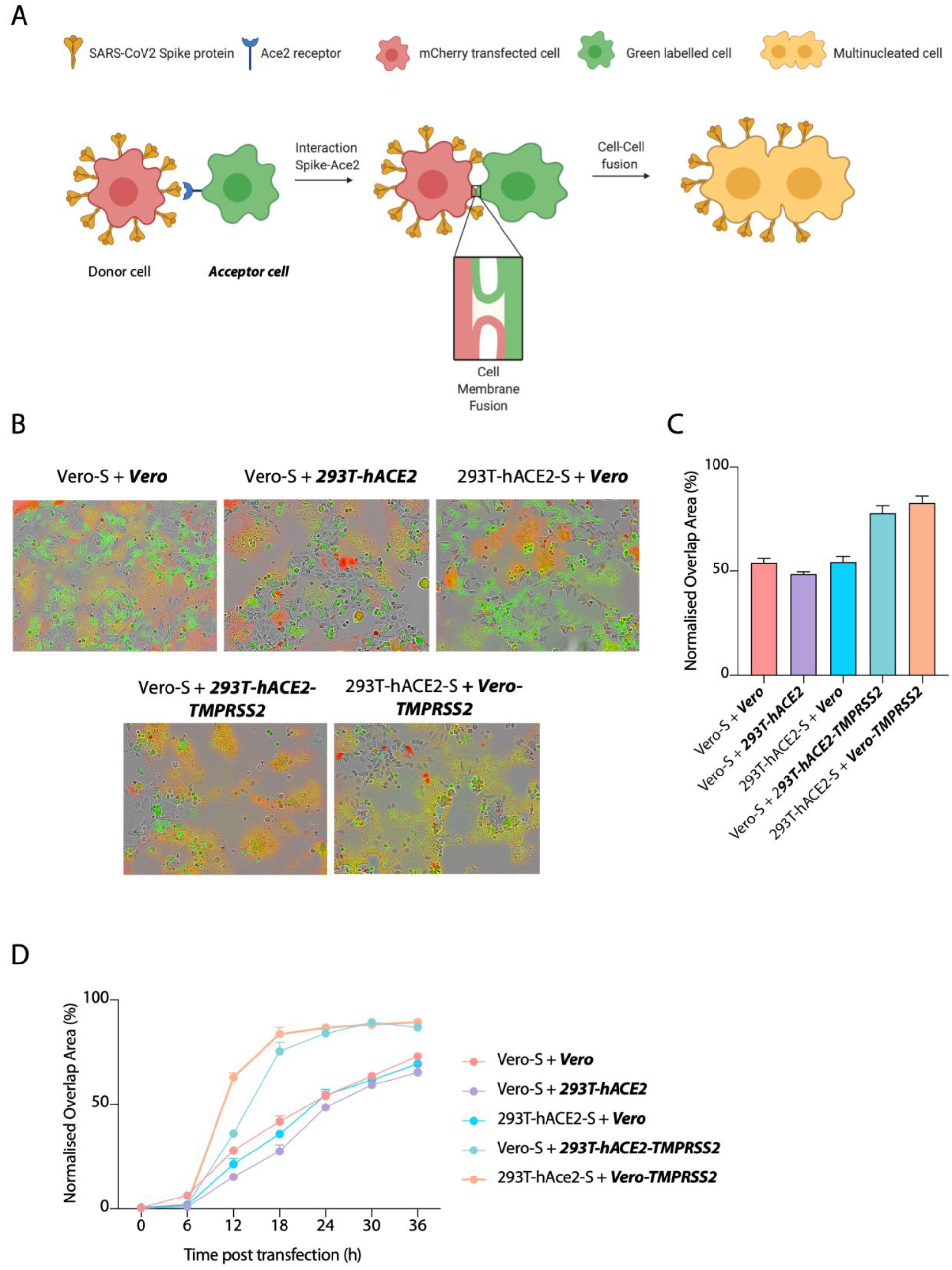
S protein overexpression induces cell-cell fusion between different cell types. **(A)** Schematic representation of S protein-mediated cell–cell fusion assay. The donor cell is identified as the cell co-expressing CoV-2 S and mCherry proteins while the acceptor cell is green-labelled. The scheme was created with BioRender. **(B)** Merged micrographs at 24 hours post transfection of the indicate cell lines transfected with CoV-2-S and mixed with green dye-labelled cells. **(C)** Quantification of (B) showing percentage of green and red overlap area at 24 hours post transfection. **(D)** Quantification of the cell-cell fusion kinetics shown in (B). Acceptor cells are marked in **bold** and *italics*.

### Furin protease enhances CoV-2 S protein cleavage but it is not required for entry in 293T cells

Furin is widely assumed to be the protease responsible for cleavage of the CoV-2 S protein multibasic site. However, most studies have relied on indirect methodologies such as small molecule inhibition or mRNA depletion (Hoffmann et al., 2020b; Shang et al., 2020; Xia et al., 2020). We decided to investigate the importance of furin directly by generating a 293T CRISPR-derived knockout cell line (herein named 293T-ΔFURIN). We confirmed that FURIN gene had been knocked out by Western blotting and as shown in Fig.S1B, there is no detectable furin protein in the knockout cells. We also generated different versions of the CoV-2 S protein harbouring amino acid substitutions in the multibasic site RRAR – either a complete inactivation by mutation to GSAS (GSAS) or a RAR amino acid deletion (ΔMBS), resembling the S protein sequence present in SARS-CoV-1 (Hoffmann et al., 2020b; Spiga et al., 2003) (Fig. 2A,B). We first analysed the role of furin and the multibasic site in S protein processing by producing lentiviruses pseudotyped with the CoV-2 S protein in both 293T-ΔFURIN cells and 293T parental cells line and analysing S that had been incorporated into viral particles. While wild-type CoV-2 S on pseudovirions produced in parental 293T cells was 80% cleaved, S protein cleavage was strongly diminished but not absent when particles were generated in the 293T-ΔFURIN cell line. This indicates that S protein processing can happen independent of furin, but that the presence of the protease strongly enhances cleavage (Fig. 2C). In contrast, pseudoviruses harbouring the GSAS mutant did not show any S cleavage when produced in either cell line, while the ΔMBS mutant showed decreased but not abolished processing (Fig. 2C). Taken together, this suggests that furin is not essential for cleavage at the S1/S2 domain boundary, but the presence of an arginine residue is. Having a multibasic sequence increases S1/S2 processing with a single arginine residue being the minimal requirement.

**Figure 2.**
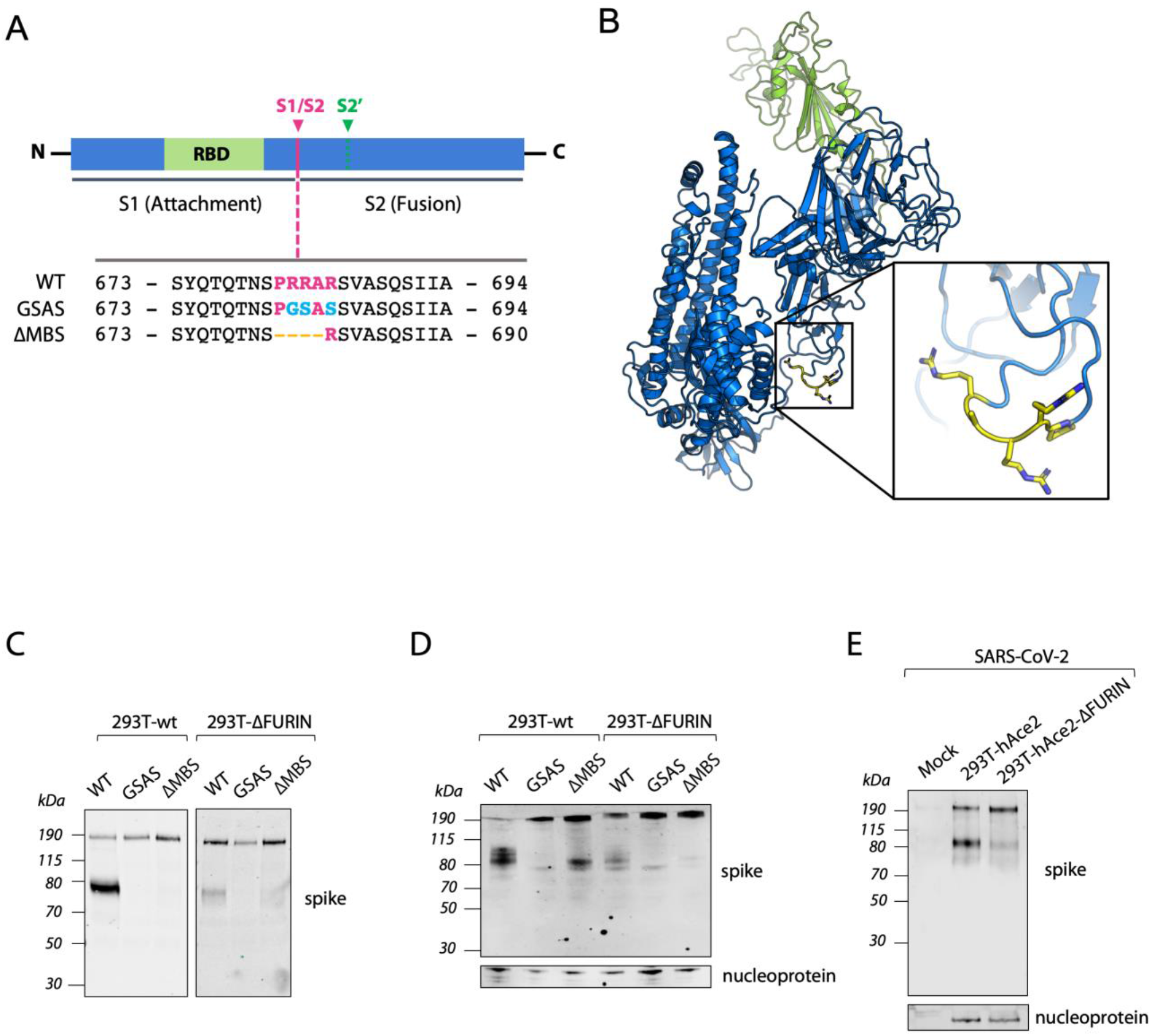
Furin is not essential for the cleavage of S protein but enhances its processing. (**A)** Schematic illustration of CoV-2 S including receptor binding domain (RBD) in green and proteolytic cleavage sites (S1/S2, S2’). Amino acid sequences around the S1/S2 recognition sites of CoV-2-S are indicated while the multibasic site is highlighted in purple. Amino acid mutations are highlighted in light blue while deletions are marked with orange dashes. **(B)** Overall structure of the CoV-2 S protein. RBD core is shown in green. Pro-Arg-Arg-Ala-Arg residues are shown in yellow. **(C,D)** A representative western blot of HIV Pseudoviruses **(C)** and Virus Like Particles (VLPs) **(D)** harbouring the indicated CoV-2 S protein mutants (detected with anti-S antibody) and produced in 293-wt and 293T-ΔFURIN cells. Expression of SARS-CoV-2 nucleoprotein is shown as loading control. **(E)** Representative western blot analysis of spike and nucleoprotein present in SARS-CoV-2 viral particles produced in 293T-hACE2 and 293T-hACE2-ΔFURIN after 42 hours post infection.

We next wanted to investigate the impact of these findings on the formation of CoV-2 virus-like particles (VLPs). In contrast to pseudotyped viruses, VLPs are entirely SARS-CoV-2 proteins, consisting of the membrane protein (M), nucleoprotein (N), envelope protein (E), and S protein. We observed that both WT and the ΔMBS mutant S proteins incorporated into CoV-2-VLPs had undergone cleavage, with almost complete processing of the WT protein and a minor fraction of the ΔMBS cleaved (Fig. 2D). Processing of VLP-incorporated S protein was furin-dependent, as the proportion of cleaved S of either WT or ΔMBS mutant was reduced in 293T-ΔFURIN cells (Fig. 2D). There was very little cleaved S protein incorporated in GSAS VLPs and this did not differ between 293T and ΔFURIN cells, suggesting that removal of all basic residues is required to fully prevent S cleavage. It also supports the hypothesis from the pseudovirus data that evolution of the multibasic site has substantially enhanced the existing S cleavage at this position, which is mediated by the single arginine site.

As a final confirmation of these results, we examined S processing in SARS-CoV-2 wild-type virus. We infected 293T-hACE2 and 293T-hACE2-ΔFURIN cell lines and collected produced virus after 42 hours. The pattern of S-cleavage observed was similar to that seen for pseudoviruses and VLPs (Fig.2E), confirming that Furin carries out, but is not essential, for S1/S2 processing. Moreover, this data proves the suitability of pseudoviruses and VLPs as surrogate models to study S protein cleavage.

Next, we sought to investigate how S cleavage at the multibasic site impacts the formation of multinucleated cells. To test this, we overexpressed CoV-2-S in 293T-ΔFURIN or 293T parental cells and measured cell-cell fusion activity. Overexpression of S in 293T-ΔFURIN cells still triggered syncytia formation when mixed with Vero cells (Fig.3A-B) although with slower kinetics compared to parental 293T cells (Fig.3C). This result is consistent with the idea that lower amounts of CoV-2-S cleavage in 293T-ΔFURIN cells is able to induce fewer cell-cell fusion events. To rule out the possibility that furin from Vero cells could cleave S produced in 293T-ΔFURIN cells, we generated a CRISPR-derived Vero cell line knocked out for furin (Vero-ΔFURIN) (Fig. S2A). Mixing the two ΔFURIN cell types, we observed that the absence of furin in acceptor cells further decreased cell-cell fusion kinetics (Fig. 3D,E), but formation of multinucleated cells still occurred (Fig. 3E, S2B), suggesting that furin from the acceptor cell contributes to syncytia formation but it is not a major determinant. Overexpression of the uncleaved GSAS mutant did not induce cell-cell fusion in any of the above-mentioned conditions suggesting an essential role for multibasic site processing in inducing syncytia (Fig. 3A-E). These data together suggest that in the absence of furin, S protein can still be cleaved by another protease that recognises the multibasic site and is able to induce the formation of multinucleated cells.

**Figure 3.**
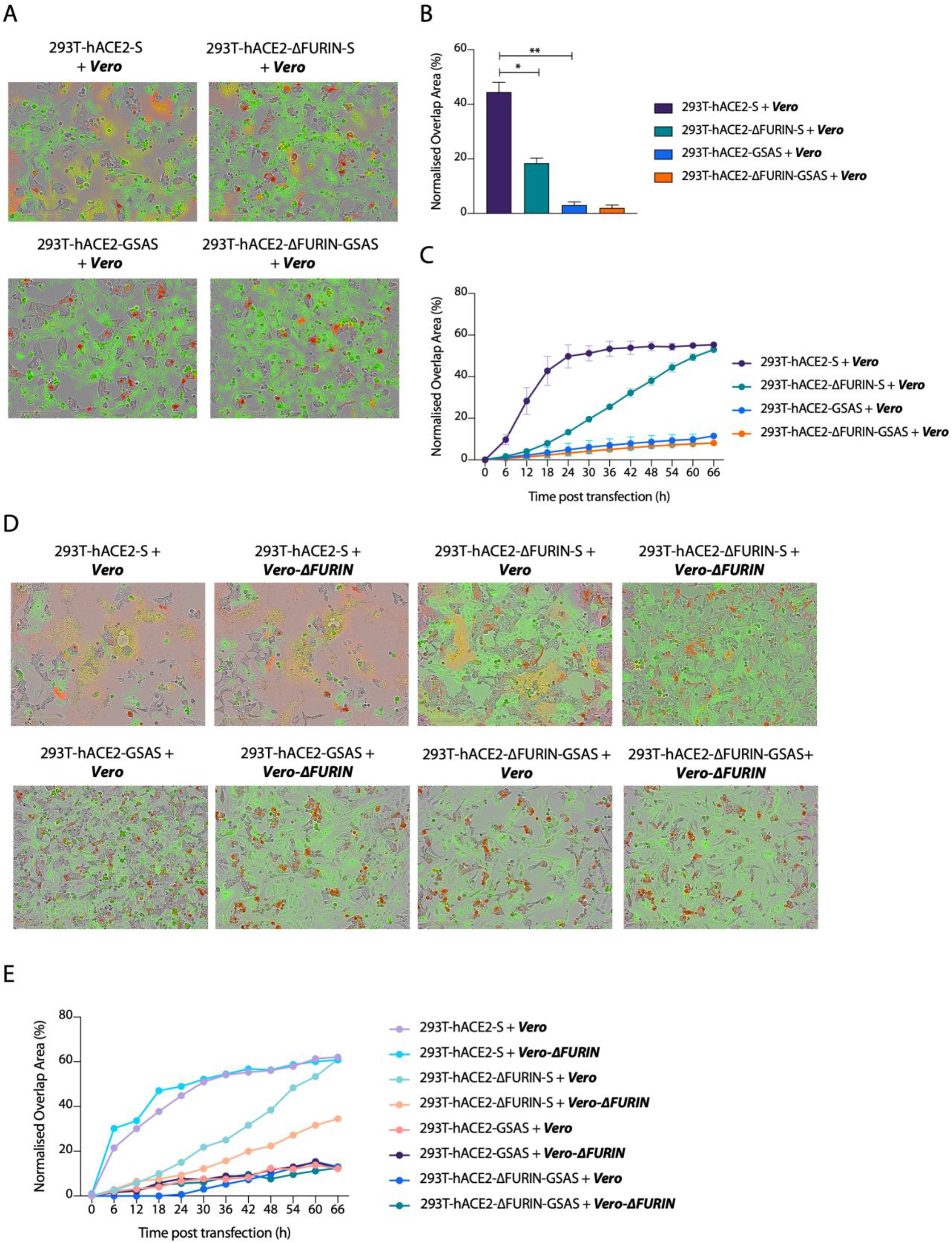
Processing of the S protein multibasic site is essential for cell-cell fusion but furin is dispensable. **(A)** and **(D)** Reconstructed micrographs of the indicated cells lines transfected with the indicated S mutants and mixed with green-labelled cells at 24 hours post transfection. **(B)** Quantification of green-red overlap area shown in (A) *P<0.05; **, P<0.001. **(C)** Kinetics of cell-cell fusion shown in (A). **(E)** Cell-cell fusion kinetics shown in (D). Data are expressed as means +/- SEM (n=2). Acceptor cells are marked in **bold** and *italics*.

### Furin ablation decreases SARS-CoV-2 replication but increases S-pseudovirus entry

Having assessed the crucial role of S cleavage in triggering cell-cell fusion we investigated how furin processing of CoV-2-S influences the infectivity of viral particles. To test this, we produced both WT and multibasic site mutants CoV-2-S pseudoviruses in 293T-ΔFURIN cells or 293T parental cells and infected 293T-hACE2 cells. Pseudoviruses harbouring wild-type S were less infectious when produced in 293T cells compared to viruses produced in 293T-ΔFURIN. This difference was lost when using pseudovirions expressing either the GSAS or ΔMBS mutant S, which increased infection compared to those expressing WT S, consistent with the fact that furin acts on the multibasic site (Fig. 4A). Many coronaviruses, including SARS-CoV-1 and SARS-CoV-2 exploit endocytosis for entry into 293T cells but the role of S cleavage in orchestrating this mechanism is not completely understood. To investigate how S processing can impact on the endocytic viral entry route, we produced pseudoviruses in 293T-ΔFURIN or 293T parental cells and compared infection of 293T-hACE2 cells in the presence of E-64d, a widely used inhibitor of lysosomal cysteine proteases. We found that S-pseudovirions produced in both ΔFURIN and parental 293T cell lines harbouring the different S mutations showed a strongly impaired entry in the presence of E-64d (Fig. 4B,C). Together these data indicate that protease-mediated S processing at the multibasic site is not required for infection of 293T cells, and that the virus is able to enter via an endocytic mechanism regardless of S cleavage.

**Figure 4.**
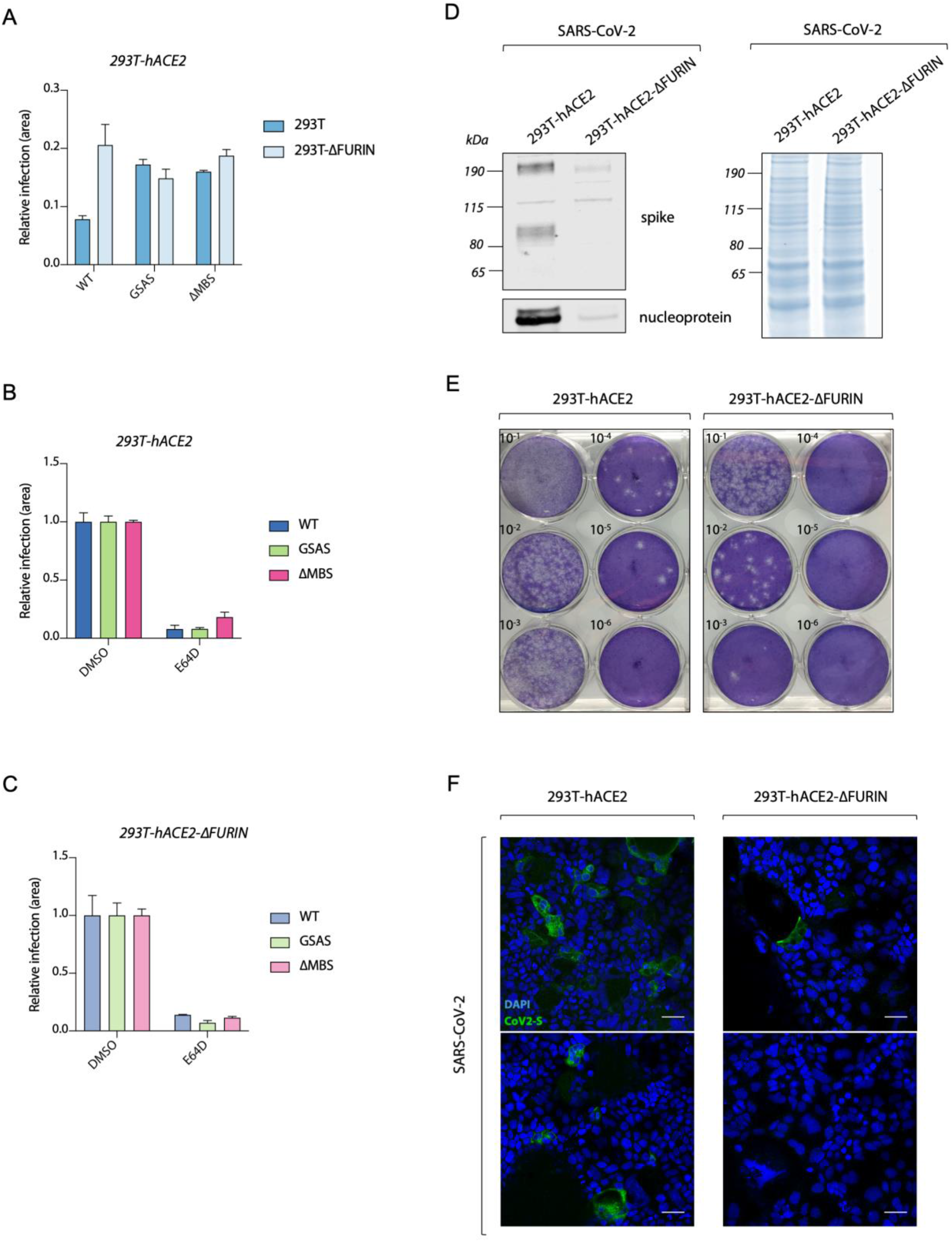
Furin enhances SARS-CoV-2 replication but is not essential for S-pseudovirus entry. **(A)** Infection of 293T-hACE2 cells with HIV pseudotyped expressing GFP and harbouring CoV-2 S mutants, measured as proportion of cell area expressing GFP. Viruses were produced in either 293T-wt or 293T-ΔFURIN cells. **(B)** and **(C)** Infection of 293T-hACE2 cells with HIV pseudotyped with CoV-2 S mutants as in (A), but pre-treated for 2 hours with either DMSO or 25μM lysosomal inhibitor E-64d as indicated. (**D)** Representative western blot analysis showing viral particles produced from SARS-CoV-2 infection of 293T-hACE2 and 293T-hACE2-ΔFURIN cells at 72 hours post infection. Spike and nucleoprotein are detected (left panel). Total protein content by Coomassie staining is shown (right panel). **(E)** Plaque assay showing infection of Vero-hACE2-TMPRSS2 with viruses produced as in D. **(F)** Immunofluorescence micrographs displaying infection of Caco2 BVDV-Npro cells with normalised amounts of SARS-CoV-2 virus produced in 293T-hACE2 and 293T-hACE2-ΔFURIN cells. Spike (green) and nuclei (blue) are shown. Scale bar, 50 μm.

Having determined that pre-cleavage of S at the multibasic site is not essential for pseudovirus entry, we sought to investigate the role of furin in SARS-CoV-2 wild-type virus replication. We infected cells with moi of 0.01 and left viruses to spread for 72 hours. Western blot analysis showed that viral particles produced in 293T-hACE2-ΔFURIN cells were markedly reduced compared to 293T-hACE2 parental cells (Fig. 4D, left panel), despite the same total protein content was present in the sample (Fig. 4D, right panel). Viral titration and plaque assay in Vero-hACE2-TMPRSS2 cells confirmed this result and showed that loss of furin leads to a reduction in the production of infectious virus by two orders of magnitude (Fig. 4E). To test whether depletion of furin reduces virion infectivity rather than numbers of virions, we normalised the amount of virus produced in 293T-ΔFURIN and parental cells by nucleoprotein expression and infected Caco2 cells, which are a model for SARS-CoV-2 infection (Chu et al., 2020). Through immunofluorescence analysis of S protein, carried out to detect infected cells at 24 hours post infection, we observed that virions produced in 293T-hACE2-ΔFURIN cells were markedly less infectious (Fig. 5F). This suggests that depletion of furin reduces the infectivity of newly made particles, probably due to the loss of S pre-processing. This result is consistent with the hypothesis that SARS-CoV-2 enters Caco2 cells mainly by virus-cell membrane fusion(Hoffmann et al., 2020a), and that pre-cleavage of S protein by furin at the S1/S2 boundary is required in order for TMPRSS2 to access and cleave the S2’ domain and trigger virus-cell membrane fusion. Thus, furin can contribute to virus spread both by increasing virion infectivity and promoting S-mediated cell-cell fusion.

**Figure 5.**
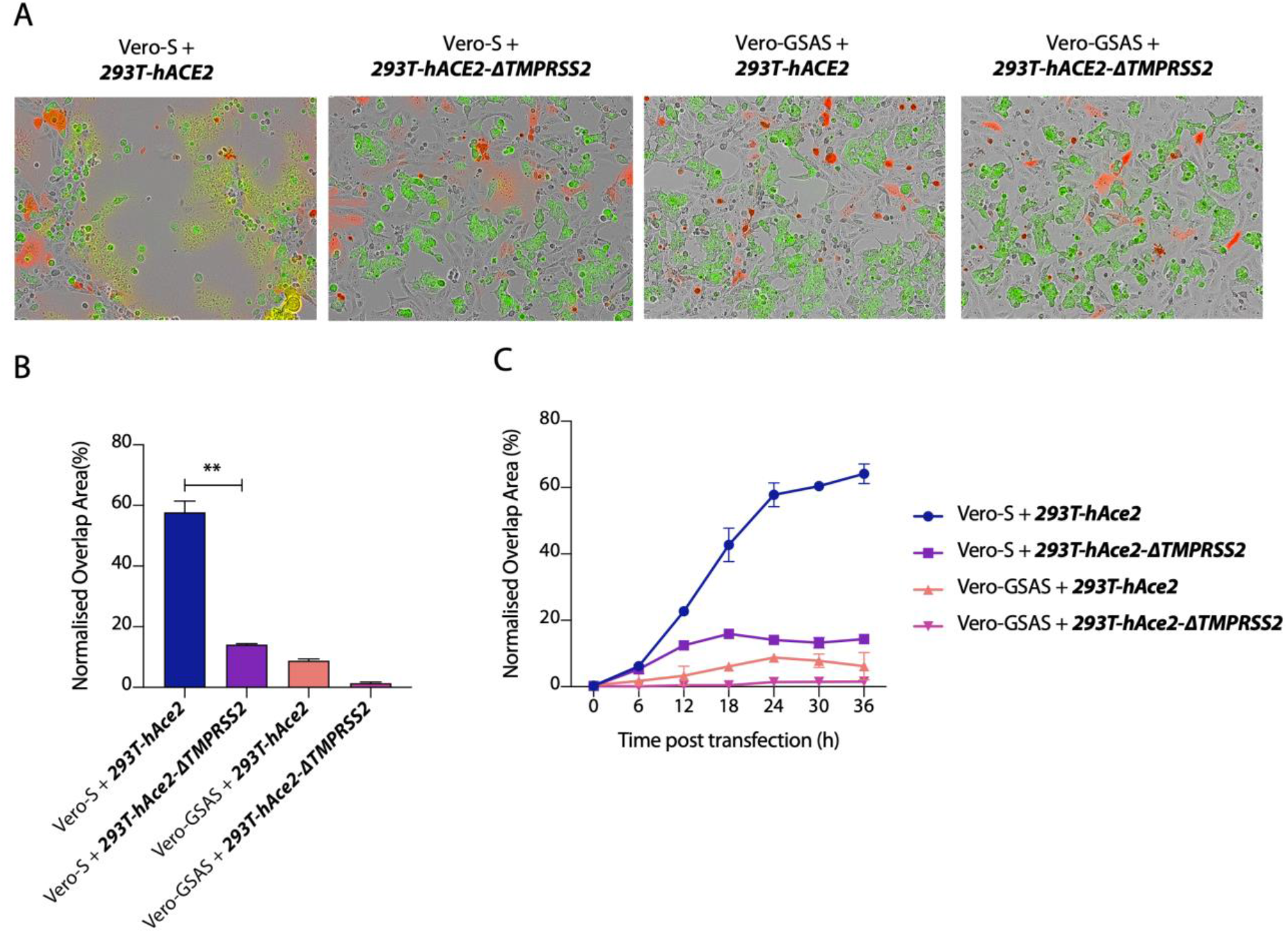
TMPRSS2 protease is required on acceptor cells to trigger cell-cell fusion. **(A)** Merged micrographs of the indicated cells lines transfected with the indicated S mutants and mixed with green-labelled cells at 24 hours post transfection. **(B)** Quantification of percentage of green and red overlap area shown in (A) at 24 hours post transfection.**P<0.001. **(C)** Quantification of cell-cell fusion kinetics as in (B). Data are expressed as means +/- SEM (n=2). Acceptor cells are marked in **bold** and *italics*.

### TMPRSS2 but not multibasic site cleavages are required for cell-cell fusion

It is thought that TMPRSS2 cleavage of the S2’ domain triggers virus-plasma membrane mixing by exposing the fusion peptide. Whether TMPRSS2 protease is similarly required for cell-cell fusion and if this favours CoV-2 virus spreading among cells, is still poorly investigated. We tackled this question with our cell-cell fusion assay and using a CRISPR-derived 293T cell line knocked out for TMPRSS2 (293T-ΔTMPRSS2) as the acceptor cell (Fig.S3A,B). The absence of TMPRSS2 protease on the acceptor cell prevented the formation of syncytia (Fig. 5A-C) showing that processing of S at the S2’ site plays a crucial role in triggering cell-cell fusion. The GSAS mutant failed to induce multinucleate cells (Fig. 5A-C) strengthening the hypothesis that S pre-cleavage at the multibasic site can facilitate the TMPRSS2-mediated processing at the S2’ site to expose the fusion peptide and trigger fusion between cell membranes.

## DISCUSSION

One of the key distinguishing features between SARS-CoV-2 and other related Coronaviruses is the presence of a multibasic site comprising several arginine residues (R) between the S1/S2 domains of the major surface glycoprotein S (Walls et al., 2020; Wrapp et al., 2020). The multibasic site has been shown to result in the cleavage of SARS-CoV-2 S (Hoffmann et al., 2020b). The protease responsible for this activity is thought to be furin but this is largely based on indirect experiments using small molecule inhibitors (Shang et al., 2020; Xia et al., 2020). Here we show that genetic knockout of furin significantly reduces but does not abolish CoV-2 S processing. This indicates that the multibasic site is also a substrate for other proprotein convertases. One candidate protease that can play a role in this process is the serine protease matriptase, which also prefers sequences with R-X-X-R amino acid sequence and has been shown to activate the hemagglutinin protein of H9N2 influenza A viruses that have the cleavage site motif R-S-S-R (Baron et al., 2013; Beaulieu et al., 2013). In the case of CoV-2 S, we found that the presence of only one R residue between the S1/S2 subunits significantly reduces cleavage efficiency and this is further reduced, but not abolished, in furin-deficient cell lines.

Importantly, our data show that furin cleavage strongly promotes SARS-CoV-2 replication. Substantially fewer virions were produced in furin knockout cells at 72 hours post infection. Plaque assays confirmed this result and showed that loss of furin decreased the production of infectious virions by two orders of magnitude. This was due to a loss of virion infectivity rather than a reduction in viral production, as when equal numbers of virions were used to infect Caco2 cells, we saw a significant reduction in S protein expression, marker of viral replication, over 24 hours. Immunoblot analysis of viruses produced in furin knockout cells showed that S protein was no longer efficiently cleaved. Taken together, this data shows that furin is required to pre-process S protein and promote infection of Caco2 cells. This is consistent with previous data suggesting that SARS-CoV-2 infects Caco2 cells via plasma membrane fusion and that pre-processing of S is required in order for TMPRSS2 to cleave S2’ and expose the fusion peptide for membrane mixing (Bestle et al., 2020; Hoffmann et al., 2020a).

Furin cleavage and the multibasic site is also thought to be important for the ability of Coronavirus S protein to mediate cell-cell fusion (Buchrieser et al., 2020; Follis et al., 2006; Xia et al., 2020). This is potentially significant for SARS-CoV-2 virulence as it provides an additional route for disseminating virus through the host. The ability to transmit between cells in a particle-independent manner also has implications for pathogenesis, as immune factors, such as antibodies, may be less able to block this type of spread. We therefore investigated whether furin is required for cell-cell fusion and the formation of multi-nucleated syncytia. Unlike with pseudovirus infection of 293T cells, we find that deletion of furin from donor 293T cells decreases S-mediated cell-cell fusion with nearby acceptor cells. Unexpectedly however, we find that while loss of furin leads to a slowing of cell-cell fusion it does not prevent it. Furthermore, we still observed syncytia formation when furin was deleted from both donor and acceptor cells. In contrast, complete removal of the multibasic site abolished S-mediated fusion. This suggests that while furin is the main protease involved in pre-processing S protein for cell-cell fusion, other proteases can cleave at the S1-S2 junction. Thus, while blocking cleavage of S1-S2 is likely to halt this mode of viral transmission, inhibitors will need to block more than just furin for complete efficacy.

A clear difference between our cell-cell fusion and pseudovirus experiments was that while pseudovirus infection and processing data are the same for GSAS mutant and ΔFURIN cells, there is a substantial variation in cell-cell spread between the two perturbations. As cell-cell fusion provided a more accurate model for furin-dependence, we used this assay to test other cofactor and cell dependencies. We found that S protein mediated cell-cell fusion was possible between different cell types, with no requirement for a particular cell type as donor or acceptor. We also found that manipulation of 293T cells greatly altered their ability to participate in fusion as overexpression of hACE2 or TMPRSS2 greatly increased syncytia formation. Conversely, knockout of TMPRSS2 from 293Ts cells abolished fusion. Fusion could not be restored when only donor cells expressed TMPRSS2. This shows that the CoV-2 S protein feature of forming multinucleated cells is dependent on the spatial orientation of TMPRSS2 in relation to the S protein. TMPRSS2 must be present in the opposing cell membrane to activate S protein and induce cell-cell fusion. This has important implications for predicting which cell types and tissues are susceptible for cell-cell viral spread: TMPRSS2 and ACE2 are essential but furin is not.

While waiting for an effective vaccine against SARS-CoV-2, antiviral drugs and specific inhibitors of enzymes involved in viral replication are urgently needed. Our findings indicate that, since furin is not essential for virus replication, inhibitors currently used against this cellular protease may be inefficient at blocking viral infection. This study also highlights the targeting of cell-cell fusion process as a potential antiviral strategy, fostering the development of specific inhibitors able to limit the spread of SARS-CoV-2.

## MATERIALS AND METHODS

### Cells

HEK293T CRL-3216, Vero were purchased from ATCC and maintained in Dulbecco’s Modified Eagle Medium (DMEM) supplemented with 10% fetal calf serum (FCS), 100 U/ml penicillin, and 100mg/ml streptomycin. All cells are regularly tested and are mycoplasma free.

Caco2 BVDV-Npro cell line was generated as described previously (Hosmillo et al., 2015).

293T-hACE2, 293T-hACE2-ΔFURIN, Vero-hACE2 were generated transducing 293T, 293T-ΔFURIN or Vero cells with lentiviral particles expressing hACE2 ORF and cultured in DMEM 10% FCS with 5 μg/ml blasticidin. 293T-hACE2-TMPRSS2 and Vero-hACE2-TMPRSS2 were instead generated transducing 293T-hACE2 and Vero-hACE2 cells respectively with lentiviral particles expressing TMPRSS2 ORF and maintained in DMEM 10% FCS with addition of 5 μg/ml blasticidin and 1 μg/ml puromycin.

### Viruses

The SARS-CoV-2 virus used in this study is the clinical isolate named “SARS-CoV-2/human/Liverpool/REMRQ0001/2020”, isolated by Lance Turtle (University of Liverpool) and David Matthews and Andrew Davidson (University of Bristol).

### Plasmids

Vectors for viral production, were, for HIV-1 Gag Pol, pCRV-1 (Naldini et al 1996) and for GFP expression, CSGW (Zennou et al 2004). Lentiviral packaging plasmid pMDG2, which encodes VSV-G envelope, was used to pseudotype infectious virions (Addgene plasmid # 12259).

For production of VLPs codon optimised plasmids encoding M,N,E,S protein were generated using DNA gblocks by IDT. The gBlocks were cloned into pcDNA3 using HindIII and XhoI restriction enzyme sites. pCAGGS-S used for cell-cell fusion assay was generated cloning codon optimised S into a pCAGGS empty backbone using EcoRI and NheI restriction sites.

pCAGGS-S Δc19 used for pseudotyping lentiviruses was generated deleting the last 19 amino acids, from the cytoplasmic tail of the S protein, which have been previously shown to contain an endoplasmic reticulum (ER)-retention signal (Ujike et al., 2016).

pCAGGS-GSAS was obtained by using QuikChange II Site-Directed Mutagenesis (Agilent Technologies). The same strategy was used to eliminate PRRA residues from pCAGGS-S harbouring the pCAGGS-ΔMBS.

The plasmids used for CRISPR-Cas9 gene editing were pSpCas9(BB)-2A-GFP (PX458, Addgene ID 48138) and pSpCas9(BB)-2A-Puro (PX459, Addgene ID 48139). pX458 and pX459 were digested with BbsI and dephosphorylated with FastAP Thermosensitive Alkaline Phosphatase. Primers containing gRNA sequences were annealed and phosphorylated by T4 Polynucleotide Kinase and the annealed oligonucleotides were inserted into the pX458 or pX459 vector by ligation. The following gRNA sequences were used: 5’ -GATGCGCACAGCCCACGTGT-3’ targeting human FURIN (in pX459, pJC142), 5’ -CGGATGCACCTCGTAGACAG-3’ and 5’ -TGTGCCAAAGCTTACAGACC-3’ targeting human TMPRSS2 (in pX459, pJC144 and pJC146 respectively) and 5’ -ATGCTTCCGTGCCACGCTAT-3’, 5’ - GCTGAGGTCCTGGTTGCTAT-3’ and 5’ -CGGTGCTATAGTGTGTATCG-3’ targeting African green monkey FURIN (in pX458, pJC228, pJC229 and pJC232).

### Virus stock

SARS-CoV-2 stock was prepared in Vero hACE2-TMPRSS2 cells. Cells were infected with 0.01 MOI of virus and incubated for 3 days. Virus stock was harvested by three freeze-thaw cycles followed by 5 min 300xg spin to remove cell debris. Titers were assessed by plaque assays in Vero hACE2/TMPRSS2 cells.

### Preparation of S pseudotyped HIV-1 virions

Replication deficient VSV-G or SARS CoV-2 pseudotyped HIV-1 virions were produced in 293T wild-type (293T-wt) cells by transfection with pMDG2 or pCAGGS-S Δc19, pCRV GagPol and CSGW as described previously (Price, Jacques et al. 2014). Viral supernatants were filtered through a 0.45μm syringe filter at 48 hours post-transfection and pelleted through a 20% sucrose cushion for 2hrs at 28000 rpm. Pelleted virions were drained and then resuspended in OptiMem.

### VLP production and S analysis

SARS-CoV-2 VLPs were produced by transfecting 2×10^6^ 293T-wt cells in T75 flasks with 25 μg total DNA using Fugene6 (Promega). The codon optimised plasmids expressing SARS-CoV-2 M, N, E, and S proteins were transfected in a ratio of 1:5:5:1. VLPs were harvested 48 h post transfection from the culture supernatants, cleared by 0.45 μm filtration, and partially purified by pelleting through a 20% sucrose layer for 2hours at 28000 rpm. VLP pellets were resuspended in 50 μl of DMEM, and the 20μl were loaded on the 4-12% Bis-Tris gel for WB analyses.

### Generation of hACE2 and TMPRSS2 lentiviral particles

For the generation of hACE2 over-expressing 293T cells, the human hACE2 ORF was PCR amplified from Addgene plasmid 1786 and C-terminally fused with the porcine teschovirus-1-derived P2A cleavage sequence (ATNFSLLKQAGDVEENPGP) followed by the blasticidin resistance gene. This continuous, single ORF expression cassette was transferred into pLenti6-Dest_neo by gateway recombination. The TMPRSS2 (gene bank accession number AF123453.1) ORF was synthesised and cloned into pLenti6-Dest_Puro by gateway recombination. Lentiviral particles were generated by co-transfection of 293T cells with pLenti6-Dest_neo_ACE2-2A-Bla or pLenti6-Dest_Puro_TMPRSS2 together with pCMVR8.74 (Addgene plasmid 22036) and pMD2.G (Addgene plasmid 12259) using PEI. Supernatant containing virus particles was harvested after 48 h, 0.45 um filtered, and used to infect cells to generate stable cell lines. Transduced cells stably expressing hACE2 or TMPRSS2 were selected with 5 μg/ml blasticidin and 1 μg/ml puromycin, respectively.

### Generation of 293T-ΔFURIN and 293T-ΔTMPRSS2

293T cells were grown to ^~^70% confluence and transfected with a mixture of 2 μg of DNA encoding the gRNA and 6 μL of Polyethylenimine (PEI, Polysciences catalogue #24765, diluted to 1 mg/mL in PBS) in 100 μL Opti-MEM. Cells were transfected with 2 μg of pJC142 to knock-out *FURIN* or co-transfected with 1 μg of pJC144 and 1 μg of pJC146 to knock-out *TMPRSS2*. 48 hours later, the cells were trypsinized and replated in complete medium containing 1.5 μg/mL puromycin. 48 hours later (day 4), cells were diluted to 1 cell per two wells in 96-well plates and grown in non-selective complete medium. Single colonies that grew were expanded and analysed by immunoblotting for *ΔFURIN* clones and genotyping PCR for *ΔTMPRSS2* clones.

For the generation of *Vero-ΔFURIN* were co-transfected with the plasmids pJC228, pJC229 and pJC232 in which a mixture containing 0.66 μg of each plasmid and 6 μL of PEI in 100 μL opti-MEM was added to cells. 48 hours later, cells were trypsinized and single GFP positive cells were sorted into each well of 96-well plates using a Synergy 1 FACS sorter. Single colonies were expanded and analysed by Immunoblotting.

### S pseudotypes infection experiments

Cells were plated into 96 well plates at a density of 7.5×10^3^ cells per well and allowed to attach overnight. Viral stocks were titrated in triplicate by addition of virus onto cells. Infection was measured through GFP expression measured by visualisation on an Incucyte Live cell imaging system (Sartorius). Infection was enumerated as GFP positive cell area. For treatment with the inhibitor E-64d, cells were pre-treated with 25μM for 2hours prior to addition of the virus.

### Cell-cell fusion assay

Acceptor cells and donor cells were seeded at 70% confluency in a 24 multiwell plate. Donor cells were co-transfected with 1.5 μg pCAGGS-S and 0,5 μg pmCherry-N1 using 6 μl of Fugene 6 following the manufacturer’s instructions (Promega). Acceptor cells were treated with CellTracker™ Green CMFDA (5-chloromethylfluorescein diacetate) (Thermo Scientific) for 30 minutes according to the manufacturer instructions. Donor cells were then detached 5 hours post transfection, mixed together with the green-labelled acceptor cells and plated in a 12 multiwell plate. Cell-cell fusion was measured using an Incucyte and determined as the proportion of merged area to green area over time. Data were then analysed using Incucyte software analysis and plotted using Prism 8 software.

### SARS-CoV-2 plaque assay

Vero hACE2-TMPRSS2 cells were seeded on 12-well plates day prior infection. Next day cells were infected with serial dilutions of the supernatant −1 to −6 for 1h and then overlayed with 0.05% agarose in 2% FBS DMEM. After 3 days cells were fixed with 4% formaldehyde and stained with 0.1% toluidine blue.

### Virus infection

293T-hACE2 or 293T-hACE2-ΔFURIN cells were infected with MOI of 0.1 or 0.01 and incubated for 42 hours or 72 hours depending on the experiment. To concentrate released virions supernatant was incubated with 10 % PEG6000 (4h at RT) and then pelleted by 30min spin at 12 000xg. Pellets were resuspended directly in Laemmli buffer with 1mM DTT. Cells were lysed with Laemmli buffer with 1mM DTT and then treated with Benzonase Nuclease(70664 Millipore) and sonicated prior loading for gel electrophoresis.

### Genotyping PCR

293T-Δ*TMPRSS2* cells were screened by genotyping PCR. Genomic DNA was extracted from wild-type and Δ*TMPRSS2* 293T cell lines using the Puregene cell kit according to the manufacturer’s instructions (Qiagen, #158388). Co-transfection of pJC144 (guide 1) and pJC146 (guide 2) was expected to result in the deletion of a large part of the *TMPRSS2* gene and in order to screen for large deletions, a pair of primers labelled as primers 1 and 3 (oJC545 5’ -GCCACCGCACCCAGCCTTGTAGTAC-3’ and oJC554 5’ -TTCCAGCAGCAGAACCACGCC-3’) were used for PCR amplification. For alleles in which large deletions had not occurred, smaller frameshift-inducing indels caused by guide 1 in the second exon were screened by PCR using the primer pair primer 1 and 2 (oJC545 and oJC552 5’ GCGACAGTGGTGTTGGGAGCAG 3’). All PCR reactions were conducted using Phusion polymerase (NEB) according to the manufacturer’s instructions with GC buffer, 1.2% DMSO, 2 ng/μl genomic DNA, an extension time of 16 seconds and an annealing temperature of 64°C. PCR products were resolved on 2.4% agarose gels with SYBR Safe (Thermo Fischer Scientific, #S33102) and relevant bands were excised from the gels, purified using the Qiagen gel extraction kit (#28704, used according to the manufacturer’s instructions) and sanger sequenced using primer 1 or 3. Alignments of sequence trace files were generated using Snapgene.

### Western Blot

Cells lines were mechanically resuspended in culture medium and centrifuged at 300 x g for 5 minutes at 4°C. Cells were washed once in PBS and resuspended in 1x LDS Sample buffer with β mercaptoethanol or in lysis buffer containing 50 mM Tris HCl pH 7.4, 150 mM NaCl, 1 mM EDTA, 0.1% Triton X-100, 1 mM PMSF (Sigma) and 1x cOmplete EDTA-free protease inhibitor cocktail (Roche). Cells were treated with Benzonase Nuclease (70664 Millipore) or were incubated on ice for a further 10 minutes prior to the clarification of the lysate at 16,100 x g for 10 minutes at 4°C respectively. Samples were then sonicated and incubated at 90°C for 5 minutes. Samples were then run on 4%–12% Bis Tris gels and transferred onto nitrocellulose membranes using an iBlot (Life Technologies).

Membranes were blocked for 1 hour in 5% non-fat milk in PBS + 0.1% Tween-20 (PBST) at room temperature with agitation, incubated in primary antibody diluted in 5% non-fat milk in PBST overnight at 4°C with agitation, washed four times in PBST for 5 minutes at room temperature with agitation and incubated in secondary antibody diluted in 5% non-fat milk in PBST for 1 hour with agitation at room temperature. Membranes were washed four times in PBST for 5 minutes at room temperature with agitation and imaged directly using a ChemiDoc MP imaging system (Bio-Rad). Alternatively, blots probed with an anti-Alexa Fluor 488 secondary antibody were imaged using a Typhoon biomolecular imager (Cytivia).

The primary antibodies used were: anti-furin (1:1000, Abcam #ab3467), anti-α tubulin (1:250, YL1/2), anti-SARS-CoV-2 S (Invitrogen, PA1-41165); anti-SARS-CoV-2 Nucleoprotein (ABIN129544). The secondary antibodies were anti-rabbit HRP conjugate (1:10000, Invitrogen 31462), anti-β actin HRP (1:5000; sc-47778), anti-rat Alexa Fluor 488 (1:3000; Thermo Fischer Scientific A-21208).

## ACKNOWLEDGEMENTS

The authors are grateful to Dean Clift and Gianluca Petris for critical reading of this manuscript. We would also like to thank John Briggs and Zunlong Ke for valuable discussions and Jia Lu for providing the Caco2 BVDV-Npro cell line.

---

**Figure S1.**
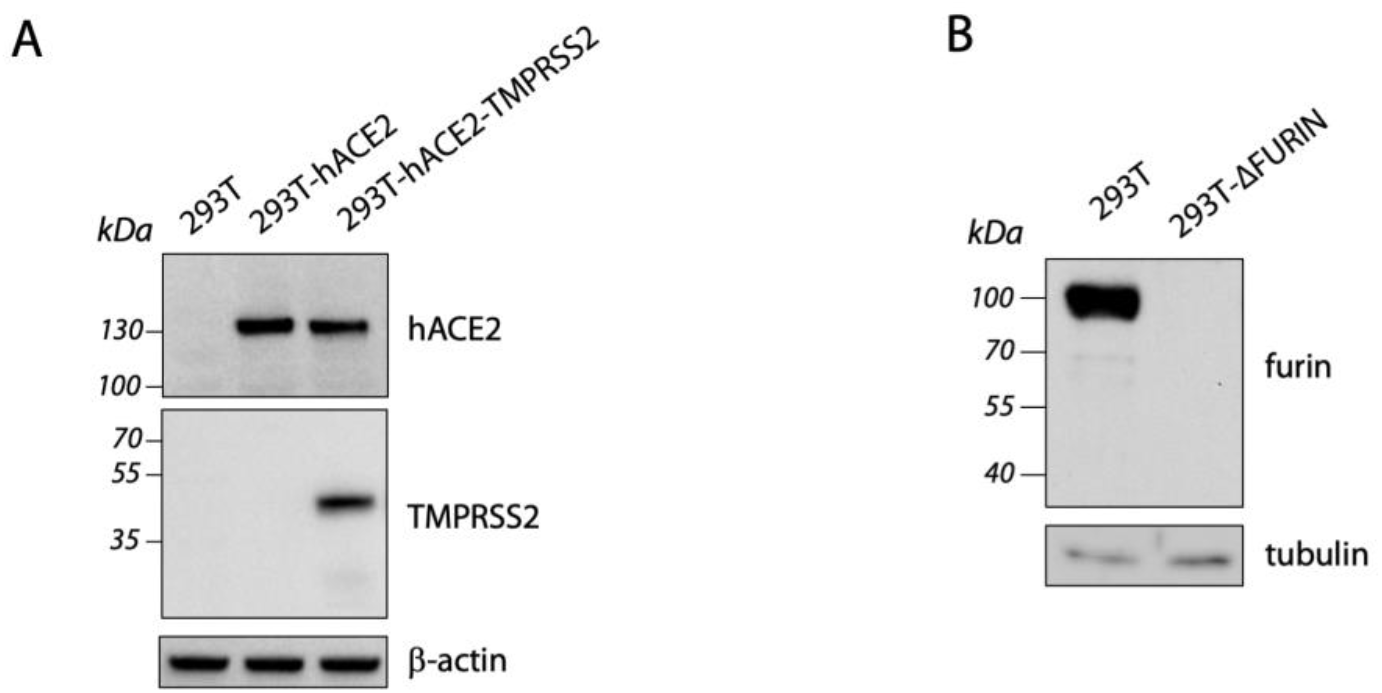
Generation of stable hACE2 and TMPRSS2 overexpressing 293T cell lines and deletion of FURIN in 293T cells by CRISPR-Cas9 gene editing. **(A)** Western blot showing hACE2 and TMPRSS2 levels in the indicated cell lines. β-actin was used as loading control. **(B)** Western blot analysis showing furin levels in 293T-ΔFURIN cell and 293T cells. Tubulin was blotted as a loading control. Immunoblots were repeated in duplicate.

**Figure S2.**
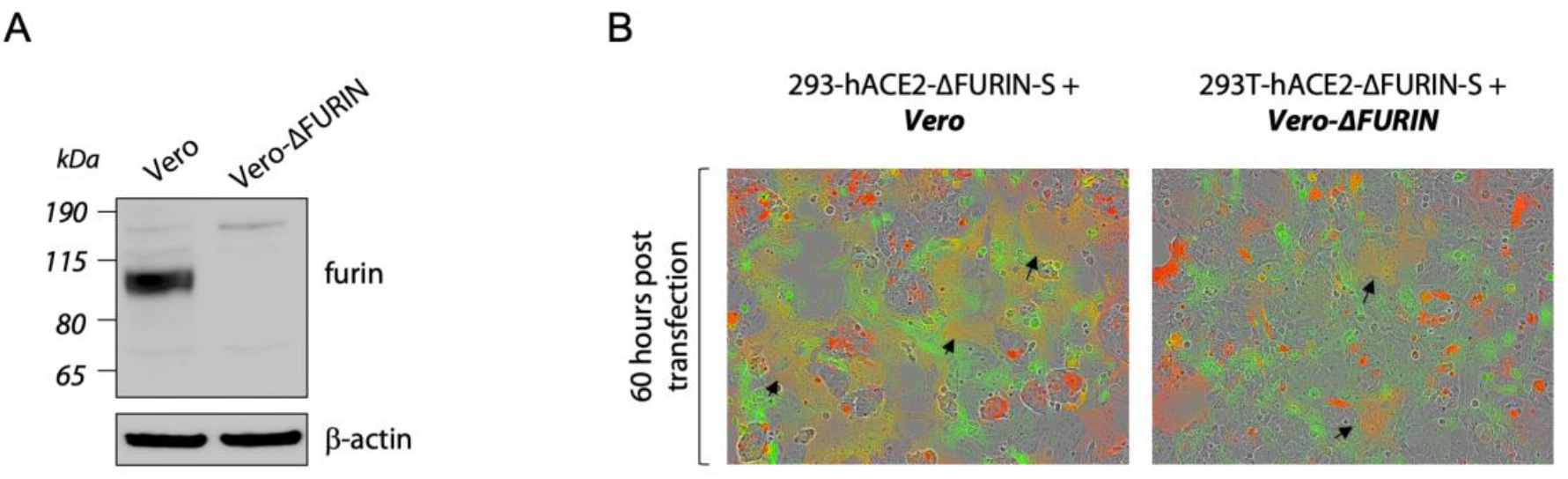
Deletion of FURIN in Vero cells did not impact the formation of multinucleated cells. **(A)** Western blot analysis showing furin levels in Vero and Vero-ΔFURIN cells. β-actin was used as a loading control. **(B)** Reconstituted micrographs of the indicated cells lines transfected with WT S and mixed with green-labelled cells at 60 hours post transfection. Black arrows indicate multinucleated cells. Acceptor cells are marked in **bold** and *italics*.

**Figure S3.**
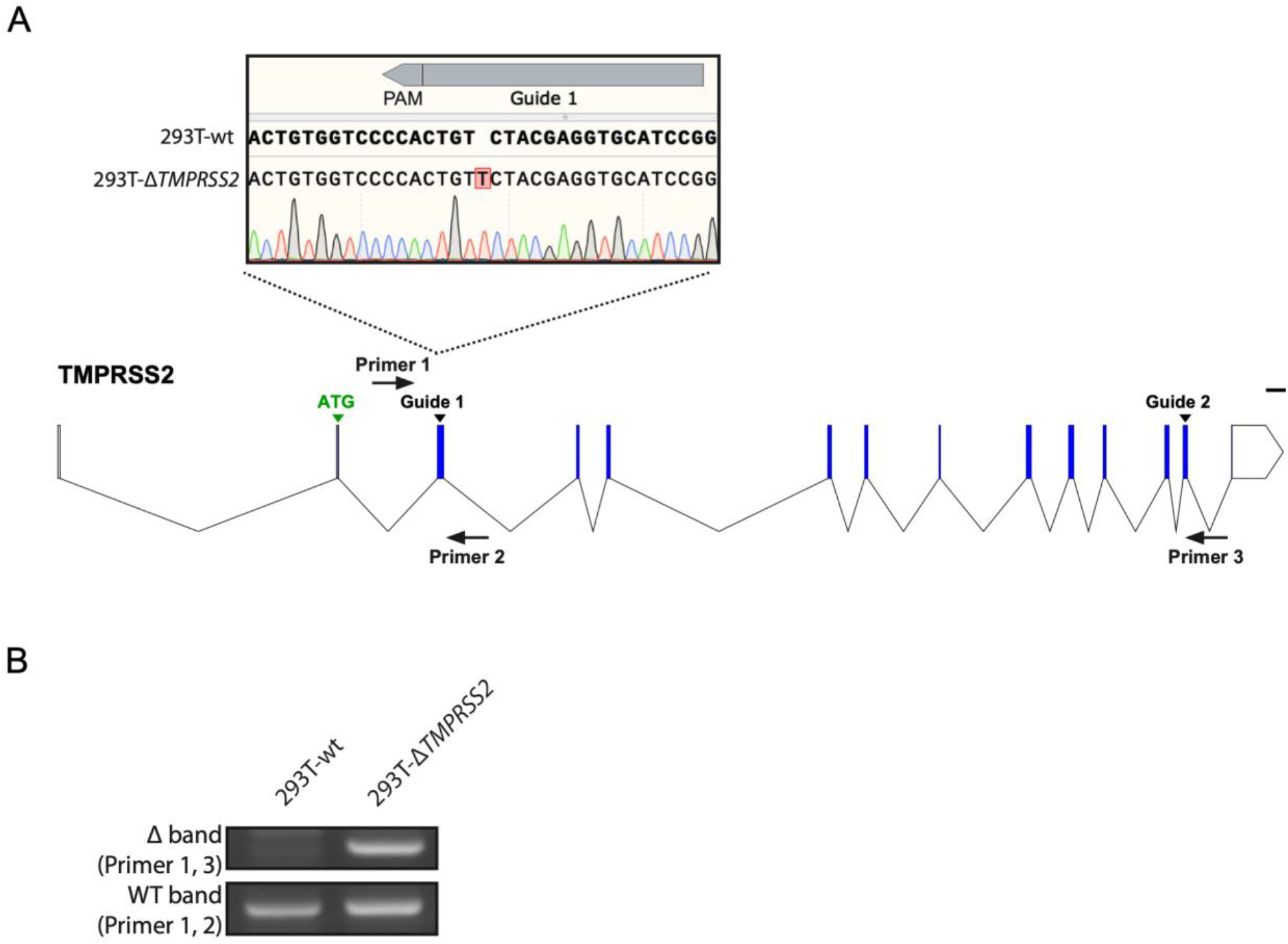
Deletion of TMPRSS2 in 293T cells by CRISPR-Cas9 gene editing. **(A)** A schematic illustrating the strategies for the deletion of TMPRSS2 and for screening deletion clones by genotyping PCR. Blue rectangles (exons), black lines (introns), scale bar = 1000 bp. Guide 1 was designed to target an early constitutive exon while Guide 2 targeted a region near the 5’UTR so as to remove a large region of the open reading frame and/or cause early frameshift-inducing indels. A pair of primers 1 and 2 was used to screen for indels by sanger sequencing (see inset) while a pair of primers 1 and 3 was used to screen for large genomic deletions by agarose gel resolution **(B)** Genotyping PCRs were conducted in duplicate.

